# A mutation associated with Charcot-Marie-Tooth disease enhances the formation of stable dynamin 2 complexes in cells

**DOI:** 10.1101/2021.12.13.472395

**Authors:** Per Niklas Hedde, Barbara Barylko, Chi-Li Chiu, Joseph P. Albanesi, David M. Jameson, Nicholas G. James

**Author notes:** Correspondence (N.G.J.); (D.M.J.); (D.M.J.).

## Abstract

Mutations in dynamin 2 (DNM2) have been associated with two distinct motor disorders, Charcot-Marie-Tooth neuropathies (CMT) and centronuclear myopathy (CNM). The majority of these mutations are clustered in the pleckstrin homology domain (PHD) which engage in intramolecular interactions that suppress dynamin self-assembly and GTPase activation. CNM mutations in the PHD interferes with these intramolecular interactions, thereby blocking the formation of the auto-inhibited state. CMT mutations are located primarily on the opposite surface of the PHD, which is specialized for lipid PIP2 binding. It has been speculated that the distinct locations and interactions of residues mutated in CMT and CNM explain why each set of mutations cause either one disease or the other, despite their close proximity within the PHD sequence. We show that at least one CMT-causing mutant, lacking residues _555_DEE_557_ (ΔDEE), displays this inability to undergo auto-inhibition as observed in CNM-linked mutants. This ΔDEE deletion mutant induces the formation of abnormally large cytoplasmic inclusions similar to those observed for CNM-linked mutant R369W. We also found substantially reduced migration from the membrane of the ΔDEE deletion mutant. These findings call into question the molecular mechanism currently believed to underlie the absence of pathogenic overlap between DNM2-dependent CMT and CNM.

## 1. Introduction

Dynamins are ∼100 kDa GTPases that catalyze membrane scission ^1^,^2^ and regulate actin polymerization^3,4^. Mammalian cells express three dynamin isoforms: dynamin 1 (DNM1), which is enriched in presynaptic nerve terminals and promotes synaptic vesicle recycling; dynamin 2 (DNM2), which is ubiquitously expressed and accounts for most examples of receptor-mediated endocytosis, and dynamin 3 (DNM3), which is enriched in postsynaptic nerve terminals and may participate in the organization of dendritic spines^5^. The three isoforms have a similar domain organization, including an N-terminal GTPase domain, a so-called “middle” domain consisting of an antiparallel three-helix bundle, a phosphoinositide-binding pleckstrin homology domain (PHD), an α-helical GTPase effector domain (GED) that folds back onto the middle domain to form the “stalk”, and a C-terminal proline/arginine-rich domain that interacts with SH3-domain-containing proteins. Dynamin self-assembly, which involves stalk-stalk interactions, stimulates GTPase activity from basal levels of ∼1-10 min^-1^ to over 200 min^-1^. Structural studies have revealed that dynamins can adopt two conformations: an assembly-competent extended (“open”) conformation, in which the PHD can bind to membranes and an auto-inhibited (“closed”) conformation, in which the PHD is folded back onto the stalk and sterically blocks self-assembly^6,7,8^.

Mutations in DNM2 have been identified in patients with two motor disorders, Charcot-Marie-Tooth disease (CMT)^9, 10^ and centronuclear myopathy (CNM)^11^, which affect nerves and muscles, respectively. There is almost no overlap between the sets of mutations that cause these two disorders, suggesting that they have distinct effects on the activity and/or regulation of DNM2. We previously reported that several CNM-linked DNM2 mutations, including R369W in the middle domain and A618T in the PHD, induce the formation of stable DNM2 polymers that are abnormally resistant to disassembly by salt and GTP^12^. Kenniston and Lemmon reported similar findings and proposed that CNM mutations disrupt allosteric regulation of DNM2 GTPase activity by the PHD^13^. They further demonstrated that the most prevalent CMT-causing mutation, K562E, did not enhance dynamin polymerization or GTPase activation but instead suppressed PIP_2_ binding and PIP_2_-stimulated GTPase activity. Subsequent structural analysis of the closed DNM1 conformation provided an explanation for these observations, as residues mutated in CNM were found to be clustered at the PHD-stalk interface, whereas residues mutated in CMT were clustered on the opposite surface of the PHD, which drives PIP_2_ binding^6,7^. Thus, CNM mutations are apparently hypermorphic due to their disruption of auto-inhibitory PHD-stalk interactions, whereas CMT mutations are apparently hypomorphic due to their disruption of activating PHD-PIP_2_ interactions. Consistent with these molecular properties, DNM2-dependent CNM and CMT are currently viewed as gain-of-function and loss-of function disorders, respectively^14,15,16^.

The direct correlation between the distinct physical and enzymatic properties of CNM- and CMT-linked DNM2 mutants and the pathogenic basis of the two diseases was challenged by our recent report showing that at least one CMT mutant (DNM2-ΔDEE, which lacks residues _555_DEE_557_ in the PIP_2_-binding surface of the PHD) shares several *in-vitro* properties with CNM-linked mutants^17^. Like the CNM mutants, DNM2-ΔDEE forms stable polymers that express high PIP_2_-independent GTPase activity *in-vitro*, and its PHD binds to PIP_2_ as strongly as the PHD of wild type DNM2 (DNM2-WT). However, we also observed that, in contrast to the CNM mutants, which form oligomers containing up to 20-25 DNM2 monomers in the cytoplasm of living cells, DNM2-ΔDEE does not self-assemble beyond the predominant dimer-tetramer equilibrium characteristic of DNM2-WT. Thus, we have extended our analysis of the behavior of DNM2-ΔDEE in live cells, focusing on the nature of its association with the plasma membrane (PM) and on its propensity to incorporate into large cytoplasmic inclusions, which are much larger than oligomers of 20-25 DNM2 monomers, which would still be of sub-diffractional size. Our data suggest that in cells, DNM2-ΔDEE displays properties intermediate between those of DNM2-WT and CNM-linked DNM2 mutants.

## 2. Results

### The CMT-associated DNM2-ΔDEE mutant forms large, stable structures on the PM of living cells

In a prior study, we found that the CNM-associated DNM2-R369W mutant forms larger and more stable complexes on the PM than DNM2-WT^18^. In Figure 1, we compare the sizes of PM puncta containing EGFP-tagged DNM2-WT, DNM2-ΔDEE, and the CNM mutant DNM2-A618T. We chose to examine the effect of the A618T mutation, which modifies the PHD, rather than the R369W mutation, which modifies the stalk, to determine if the formation of large PM complexes is a common feature of CNM-linked mutations. Total internal reflection (TIRF) microscopy imaging (Fig. 1) of transfected cells clearly demonstrated that both DNM2-ΔDEE and DNM2-A618T form larger PM puncta than DNM2-WT.

**Figure 1.**
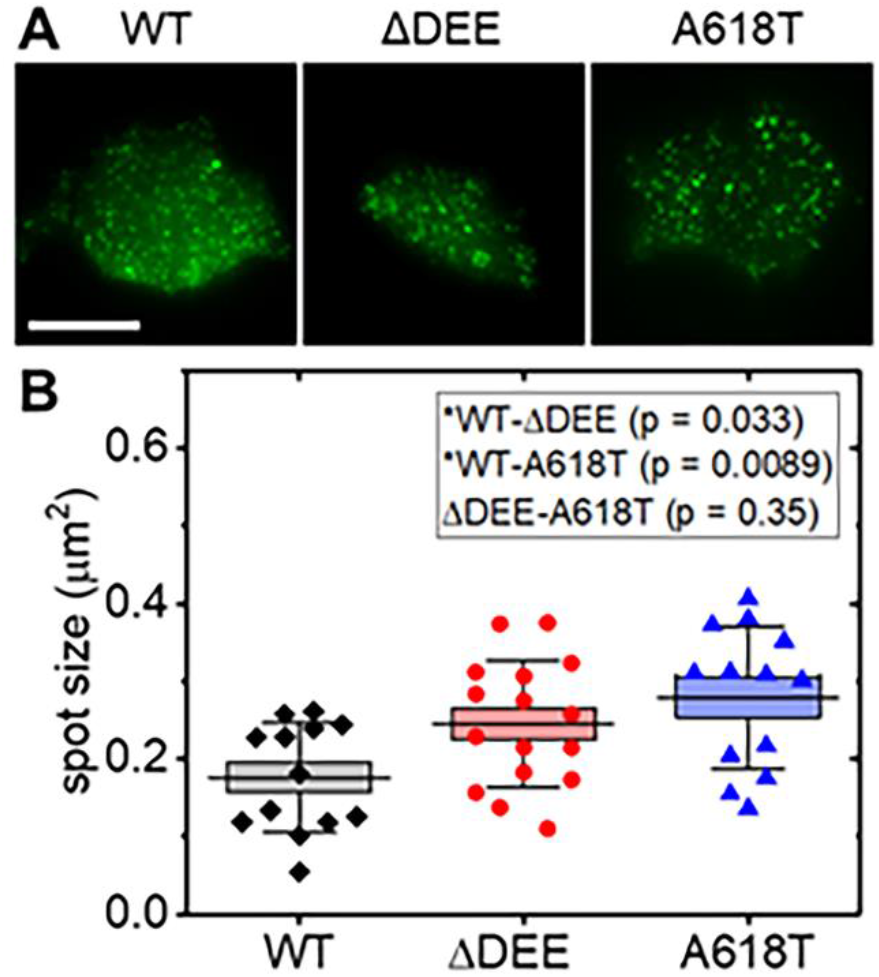
Size comparison of DNM2-containing plasma membrane inclusions in HEK293T cells expressing EGFP-tagged DNM2 constructs. (A) Representative TIRF images of cells transfected with DNM2WT-EGFP (left), DNM2-ΔDEE-EGFP (middle), and DNM2-A618T-EGFP (right). Scale bar, 20 μm. (B) Average area occupied by the inclusions. Black squares: DNM2WT; red circles: DNM2-ΔDEE; blue triangles: DNM2-A618T. Statistical analysis confirms that membrane inclusion sizes for DNM2-ΔDEE and DNM2-A618T are significantly larger than for DNM2-WT. P values (insets) were calculated with the Mann-Whitney non-parametric test; box: standard error; whiskers: standard deviation; center line: mean.

The dynamics of PM puncta containing fluorescent DNM2 were then examined using three-dimensional single particle tracking (3D-SPT)^19^,^20^. Figure 2A shows a representative 3D-SPT time trace for DNM2-WT, wherein the tracked membrane punctum (t = 0 s, highlighted in blue) was internalized in the axial direction shortly after tracking was initiated. Fluorescence was no longer detectable once the punctum had traveled ∼60 nm deep into the cytoplasm (highlighted in red). Loss of fluorescence is likely due to the disassembly of DNM2-WT polymers following internalization, as low-order DNM2 oligomers would not be detectable in our tracking setup. All of the DNM2-WT-containing puncta reported in this work were found to track several nanometers in the axial direction (61 ± 31 nm), and their signals could no longer be tracked after about one minute (80 ± 50 s). We note that the large variance in both distance and acquisition time likely results from initiating data collection at arbitrary time points within the life cycle of the membrane puncta. In Figure 2B, we show the behavior of one DNM2-WT-containing punctum and one DNM2-ΔDEE-containing punctum, which was typical of all puncta examined. In contrast to the DNM2-WT punctum, the punctum containing DNM2-ΔDEE hovered within about 10 nm of the plasma membrane, showing no sign of internalization; tracking was eventually stopped after 15 min. Figure 2C relates the distances traveled by each punctum as a function of the time at which fluorescence became undetectable (for DNM2-WT) or at which tracking was terminated (for DNM2-ΔDEE and DNM2-R369W). Puncta containing DNM2-ΔDEE or DNM2-R369W showed minimal changes within the axial direction (15 ± 10 nm), even with observation times extending to almost 20 minutes. Taken together, results shown in Figures 1 and 2 demonstrate that DNM2-ΔDEE forms large, stable structures on the PM of live cells, thus revealing another similarity with the CNM-linked DNM2 mutants.

**Figure 2.**
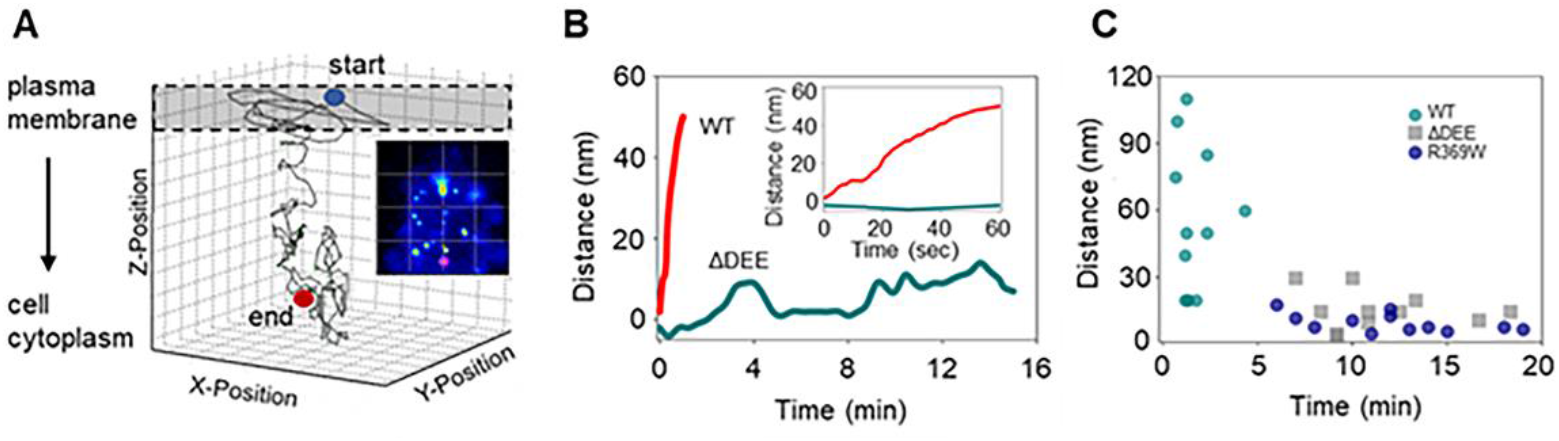
3D-SPT of DNM2 membrane puncta in HEK293T cells. (A) Representative 3D-SPT time trace for DNM2-WT with the start and end points of the time trace marked in blue and red, respectively. The shaded area represents the plasma membrane. The tracked DNM2WT membrane punctum was internalized shortly after tracking was initiated and no longer detectable once it had traveled ∼60 nm deep into the cytoplasm. The gray box represents the membrane region. (B) Exemplary time traces of one DNM2-WT-containing (red track) and one DNM2-ΔDEE-containing punctum (cyan track). For the punctum containing DNM2-ΔDEE, which hovered within ∼10 nm of the plasma membrane showing no sign of internalization, tracking was stopped after 15 min. (C) Distances traveled by 11 (DNM2-WT) 10 (DNM2-ΔDEE), and 12 (DNM2-R369W) tracked puncta as a function of the time. Tracking was stopped if fluorescence became undetectable (for DNM2-WT) or after 5-20 minutes (for DNM2-ΔDEE and DNM2-R369W). Puncta containing DNM2-ΔDEE or DNM2R369W showed minimal changes in axial direction (15 ± 10 nm), even for observation times over ten minutes.

### The ΔDEE mutation enhances the formation of cytoplasmic DNM2-containing puncta

We previously reported that the CNM-associated R369W mutation increases the propensity of DNM2 to incorporate into large puncta in the cell cytoplasm^18^. A similar observation was subsequently reported for the CNM-associated S619L mutation^*8*^. Although the clinical significance of these puncta remains to be defined, similar structures have been identified in biopsies from CNM patients and in animal models of CNM (see Discussion). In light of our findings that the CMT-associated ΔDEE mutation confers gain-of-function properties to DNM2 that had previously only been observed in CNM-linked mutants, we carried out a quantitative comparison of cytoplasmic puncta formed by DNM2-WT, DNM2-ΔDEE, and the CNM-linked mutant DNM2-A618T. Figure 3A shows representative confocal images of NIH-3T3 cells transfected with EGFP-tagged DNM2-WT, DNM2-ΔDEE, and DNM2-A618T at low, medium, and high expression levels. Visual inspection of these images demonstrates that both DNM2-A618T and DNM2-ΔDEE have a higher propensity to form large cytoplasmic puncta than DNM2-WT. Figure 3B displays the sizes of these puncta as a function of the average fluorescence intensity of the cell, which is proportional to the DNM2 expression level. Statistical analysis of these data confirmed that at low expression levels, both DNM2 mutants tend to form larger puncta than DNM2-WT (Figure 3C). At high expression levels, DNM2-WT can also form large puncta, although to a significantly lower extent than DNM2-A618T (Figure 3D). These results indicate that DNM2-ΔDEE forms cytoplasmic puncta that are intermediate in size between those formed by DNM2-WT and DNM2-A618T.

**Figure 3.**
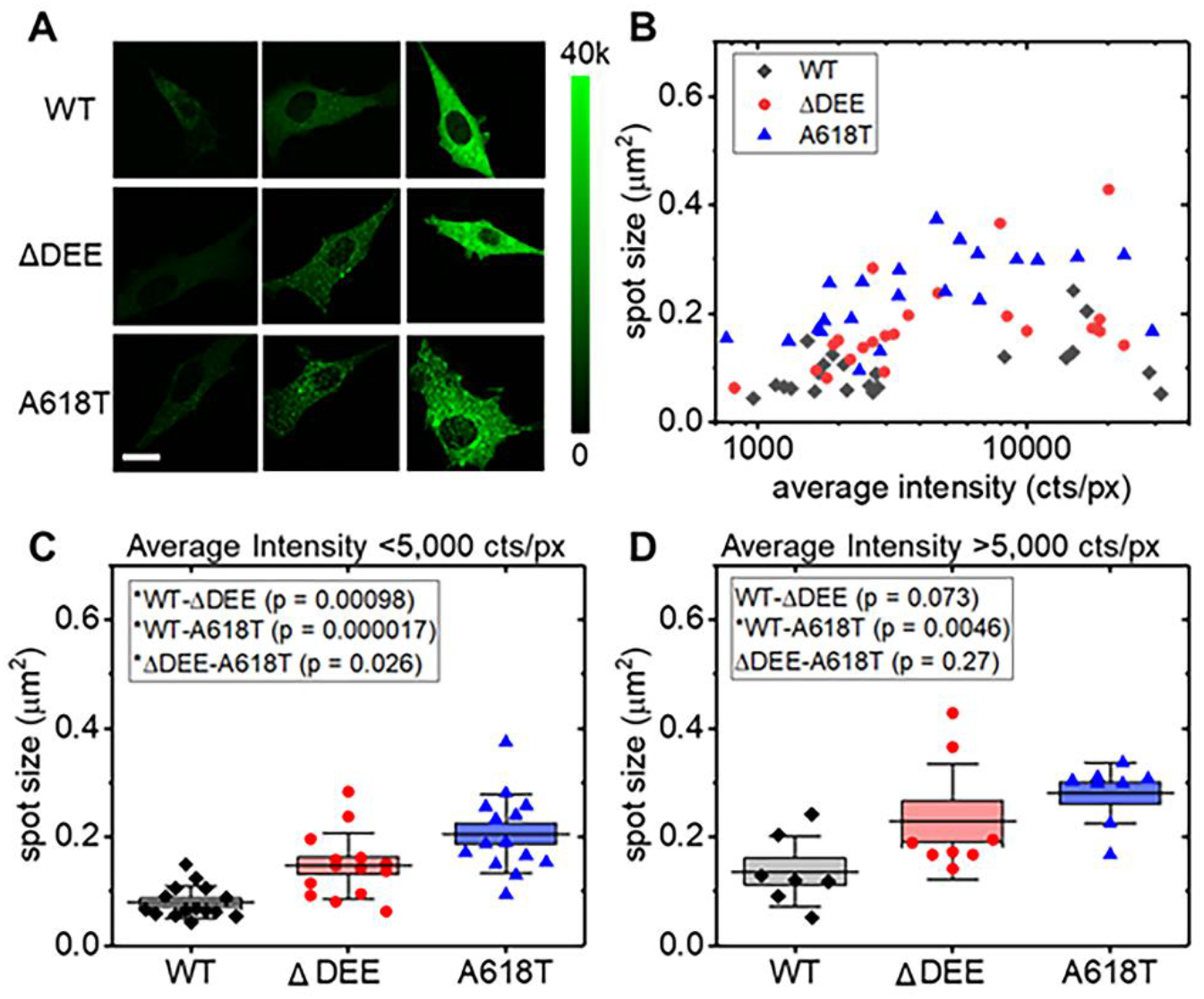
Concentration-dependent size comparison of DNM2-containing cytoplasmic inclusions in NIH-3T3 cells expressing EGFP-tagged DNM2 constructs. (A) Representative confocal images of low- (left column), medium- (center column), and high-expressing cells (right column) transfected with DNM2-WT-EGFP (top row), DNM2-ΔDEE-EGFP (center row), and DNM2-A618T-EGFP (bottom row). Scale bar, 20 μm. (B) Average area occupied by the inclusions as a function of the average intensity per pixel proportional to the protein expression level. Black squares, DNM2-WT; red circles, DNM2-ΔDEE; blue triangles, DNM2-A618T. The graph indicates that, at low expression levels, aggregate formation is low for DNM2-WT, but high for both DNM2-ΔDEE and DNM2-A618T. At high expression levels, DNM2-WT also tends to form inclusions. Each data point represents the average spot size obtained in a single cell. (C) Statistical analysis confirms that inclusion sizes for DNM2-ΔDEE and DNM2-A618T are significantly larger than for DNM2-WT for low-expressing cells (<5,000 cts/px). (D) For high expressing cells (>10,000 cts/px), only DNM2-A618T shows a significant increase in spot area over DNM2-WT. P values (insets) were calculated with the Mann-Whitney non-parametric test; box: standard error; whiskers: standard deviation; center line: mean.

### DNM2-containing cytoplasmic puncta are membrane-less structures

We next employed the technique of fluorescence recovery after photobleaching (FRAP) to determine whether the DNM2-containing cytoplasmic puncta represent membrane-enclosed organelles, such as secretory or endocytic compartments, or are more similar to membrane-less biomolecular condensates. FRAP allowed us to investigate the exchange rate of proteins within those cytoplasmic inclusions with the surrounding environment. Figure 4 shows pre-bleaching (panels A and B) and post-bleaching (panels C and D) fluorescence images of DNM-WT (panels A and C) and DNM2-ΔDEE (panels B and D). The photobleached puncta are marked by red circles. The average normalized fluorescence recovery data of DNM2-WT and DNM2-ΔDEE (10 cells each) are displayed in Figure 4E. Statistical analysis of the recovery times obtained by fitting a single exponential association model did not reveal any significant difference between DNM2-WT and DNM2-ΔDEE, despite the difference in average sizes of the inclusions formed by these proteins (Figure 3). These results demonstrate that the cytoplasmic inclusions formed by DNM2 are membrane-less structures and that the DNM2 molecules within them exchange rapidly with their counterparts in the surrounding environment.

**Figure 4.**
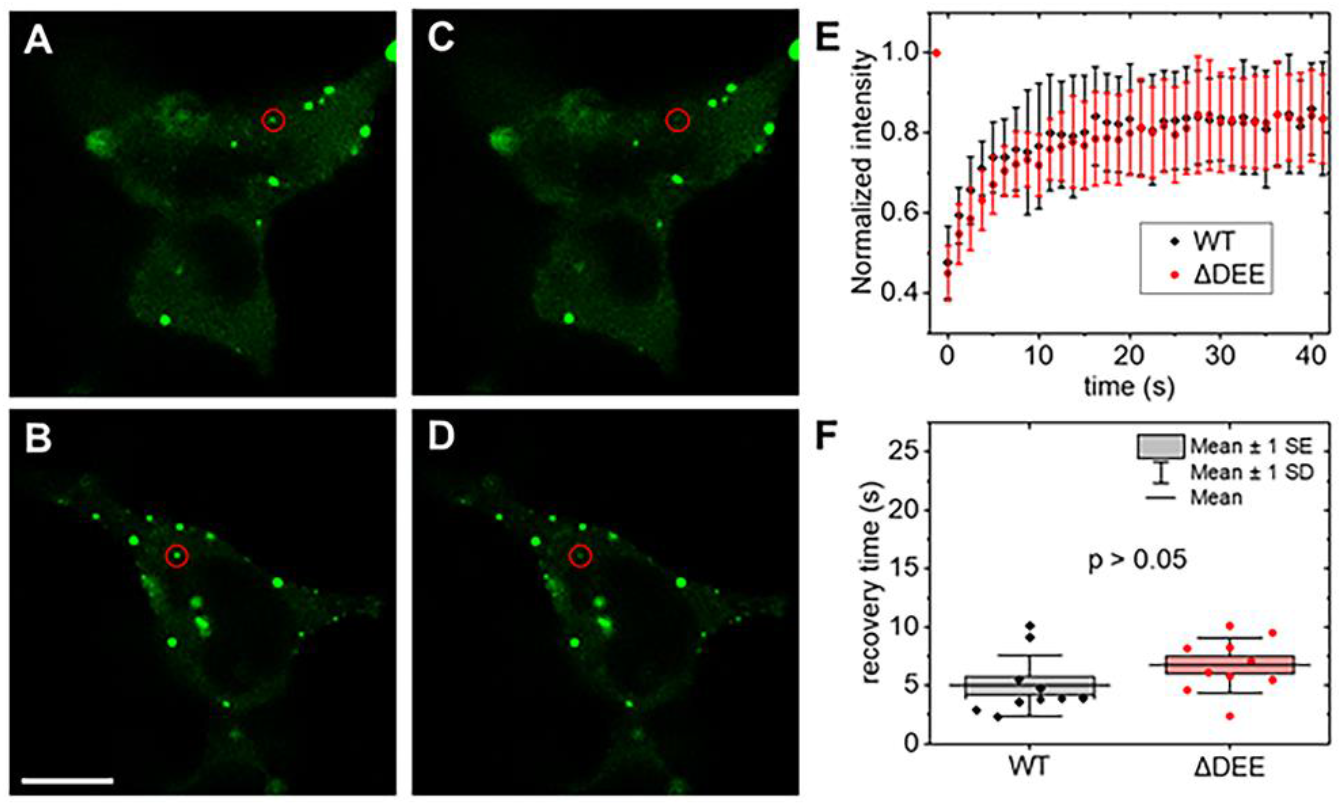
FRAP of cytoplasmic DNM2 inclusions in HEK293T cells. (A,B) Pre-bleaching and (C,D) post-bleaching fluorescence images of DNM2-WT-EGFP (A,C) and DNM2-ΔDEE-EGFP (B,D). Bleached inclusions are marked by red circles. (E) Average normalized fluorescence recovery data of DNM2-WT-EGFP (black diamonds) and DNM2-ΔDEE-EGFP (red dots) of N = 10 cells each. (F) Statistical analysis of recovery times obtained by fitting a single exponential association model to each data set. The p value was calculated by the non-parametric Mann-Whitney test. Outliers were excluded according to the Grubbs test. Scale bar, 10 μm.

## 3. Discussion

The molecular basis of dynamin-dependent motor disorders remains unclear. Therefore, it is important to characterize the dynamin mutants that cause these disorders *in-vitro* and in cells^14^. All CNM-associated DNM2 mutants examined to date have displayed enhanced abilities to self-assemble *in-vitro*^12, 13^ and, as a result of their enhanced self-assembly, increased levels of PIP_2_-independent GTPase activation^13^. These gain-of-function properties are consistent with the observed effects of expression of these mutants in cells, animal models, and patient samples. For example, overexpression of the CNM-associated mutants R465W, A618T, and S619L (but not of the G537K CMT-linked mutant), induced fragmentation of T-tubules in Drosophila muscle cells. CNM-like histopathological phenotypes were also observed in mice^15^,^21^ and zebrafish^22^,^23^ expression of CNM-associated DNM2 mutants. The gain-of-function nature of DNM2-dependent CNM is further highlighted by findings that CNM-like phenotypes were induced upon increased expression of DNM2-WT in mice^24^,^25^, whereas these phenotypes were reversed upon reduction of DNM2 expression^26^. Notably, reduction of DNM2 expression caused impairment of myelination, a characteristic of DNM-dependent CMT^27^.

In contrast to the enhancement of DNM2 self-assembly and GTPase activation induced by CNM-causing mutations, the major effect of the best-characterized CMT mutation, K562E, is near-total suppression of DNM2 interaction with PIP_2_, resulting in its inability to express PIP_2_-stimulated GTPase activation ^13^. Because PIP_2_ binding is essential for dynamin function in clathrin-mediated endocytosis (CME)^28^,^29^,^30^, DNM2-K562E acts as a dominant-negative inhibitor of CME in cells ^13^. Although residues _555_DEE_557_ are in close proximity to K562 and the PIP_2_-interaction loops of the PHD, their deletion has no effect on its binding to PC/PIP_2_-containing vesicles^17^ or on CME^31^,^32^,^33^. Instead, DNM2-ΔDEE expression has been found to impair clathrin-independent endocytosis,^33^ and to cause dynamic instability of microtubules^31^.

Our findings that DNM2-ΔDEE forms large, stable complexes on the PM and incorporates into large membrane-less cytoplasmic inclusions are consistent with our recently reported *in-vitro* results indicating that, like the CNM mutants, DNM2-ΔDEE polymers are more resistant to disassembly than DNM2-WT polymers^17^. Impaired disassembly of CNM-associated DNM2 mutants was explained by structural investigations demonstrating that the auto-inhibitory stalk-PHD interaction is inhibited in these mutants^6,7,8,34^. This explanation may not apply to DNM2-ΔDEE, as residues _555_DEE_557_ are located on the opposite surface of the PHD and, hence, would not be expected to disrupt the stalk-PHD interface. Current work is directed at testing whether the ΔDEE mutation affects the orientation of the C-terminal PRD, which is a positive regulator of dynamin self-assembly^35^ and a determinant of the different propensities of DNM1 and DNM2 to polymerize^36^.

At present, the composition of the DNM2-containing membrane-less inclusions and their contribution to disease remain to be established. However, aggregate-like DNM2 puncta were previously observed in muscle biopsies from a patient expressing the CNM mutant DNM2-D614N^37^ and that large intracellular structures containing the CNM-associated DNM2-R465W mutant, expressed at approximately endogenous levels, were observed in knock-in mice^21^. For DNM2-R369W, we also noted the formation of large cytoplasmic inclusions in CV1 cells18 and, most recently, several other novel CNM-associated DNM2 mutants (G495R, V520G, G624V, P294L, and R724H) have been associated with the formation of cytoplasmic inclusions in C2C12 cells^38^. DNM2-R465W, which causes a mild-to-moderate form of CNM, was shown to form small aggregates along the sarcolemma of transgenic zebrafish, whereas DNM2-S619L, found in more severe cases of CNM, formed numerous large aggregates located throughout the sarcoplasm^23^. Notably, the CMT-linked mutant DNM2-G537C did not form aggregates in transgenic zebrafish.

## 4. Materials and Methods

### Cell culture

HEK293 and NIH-3T3 cells were cultured in DMEM (supplemented with 10% FBS and 1% Pen-Strep), plated on fibronectin-coated glass-bottom dishes (No 1.5), and transfected with Lipofectamine 3000 (ThermoFisher, Waltham, MA). Cells were imaged 18-28 h after transfection.

### Constructs

Each fluorescent DNM2 construct utilized was cloned into the pEGFP-N1 vector (Clontech) as described in Tassin et al.^*18*^ to generate C-terminal EGFP fusion constructs.

### Confocal imaging

NIH-3T3 cells were transfected with plasmids encoding either DNM2-WT, DNM2-ΔDEE, or DNM2-A618T and imaged at 37ºC and 5% CO2 with a LSM880 laser scanning microscope (Zeiss, Jena, Germany). EGFP fluorescence was excited at 488 nm and collected in a band of 510-560 nm with a 40x, NA 1.2 water immersion lens and focused on a pinhole set to one Airy unit before detection with a photomultiplier detector. Images of 1,024 × 1,024 pixels were acquired with a pixel size of 83 nm. All imaging parameters were kept constant during the experiment such that photon counts in each pixel were proportional to the fluorescent protein levels.

### Fluorescence recovery after photobleaching (FRAP)

HEK293T cells were transfected with plasmids encoding DNM2-WT and DNM2-ΔDEE and imaged at 37ºC and 5% CO2 with a LSM880 laser scanning microscope (Zeiss) set up for FRAP. EGFP fluorescence was excited at 488 nm and detected in a band of 510-560 nm with a 40x, NA 1.2 water immersion lens. In each data set, one pre-bleaching and 34 post-bleaching frames of 256 × 256 pixels were acquired at 1.25 s intervals with a pixel size of 138 nm. EGFP was bleached in 1-3 circular regions per cell of 1-2 μm diameter for 0.3-1 s each using 100% laser power. Data were analyzed in Origin (OriginLab, MA). A single exponential association model was used to calculate the recovery times for each experiment.

### Total internal reflection microscopy (TIRFM)

HEK293T cells were transfected with plasmids encoding either DNM2-WT, DNM2-ΔDEE, or DNM2-A618T and imaged at room temperature with an Olympus IX81 (Olympus) equipped with a TIRF illumination module. EGFP fluorescence was excited with 488-nm light at the critical angle, collected in a band of 510-540 nm with a 60x, NA 1.45 oil immersion lens and, after an additional 2x magnification (total magnification 120x), imaged with a Prime95B (Photometrics) sCMOS camera. Per image, 10 frames were acquired with 100 ms exposure time and averaged before analysis.

### Spot size quantification

To quantify spot sizes in the resulting image data, a custom Matlab (R2019a, MathWorks, Natick, MA) script was used. In each image, the cell outline was traced by the user to create a mask for analysis. In this region, the image was median-filtered in windows of 3 pixels to create a de-noised image (med3) and 25 pixels to create a low-pass filtered image (med25). In addition, the detector background (bgk) was determined in a region not showing any fluorescence. A binary mask (bm) was created by subtracting 1.5 times the low frequency image and background from the de-noised image (bm = med3 – 1.5 * med25 – bkg) followed by application of a zero-intensity threshold (bm (bm <0) = 0). Individual spots were then identified as concatenated areas of pixels with more than eight neighboring non-zero pixels (Matlab function, bwconncomp(bm,8)).

### 3D orbital tracking

The hardware and software for the three-dimensional single particle tracking (3D-SPT) routine were implemented on a FV1000 (Olympus, Center Valley, PA) confocal microscope as previously described^*19*^. The galvano scanner and z-nano-drive of the microscope were driven by an IOtech DAC card (MCC, Norton, MA) to control the laser beam position. HEK293 cells, which were plated onto fibronectin-coated No 1.5 coverglass imaging dishes, were transfected with DNM2 constructs using Lipofectamine 2000. Cells were imaged with a 40x 0.8 NA water immersion objective with a working distance of 3 mm. The argon-ion laser of the Olympus FV1000 was set to an excitation wavelength of 488 nm (0.1% power) to excite EGFP and the emission was collected in a band between 505 nm and 605 nm. The fluorescence intensity at 128 points along a circular orbit was collected with a pixel dwell time of 128 μs (16.4 ms/orbit). All tracking data were analyzed using SimFCS software (available at: https://www.lfd.uci.edu/globals/).

## 5. Conclusions

In conclusion, the present study indicates that the gain-of-function properties of DNM2-ΔDEE that we had observed *in-vitro*^17^ may also be manifested in live cells. Given the similarities between DNM2-ΔDEE and the CNM-associated DNM2 mutants, it is unclear why the ΔDEE mutation has only been identified with CMT neuropathies and raises the larger question of the molecular basis underlying the lack of pathogenic overlap between DNM2-dependent CMT and CNM.

## Author Contributions

All authors have contributed to writing the manuscript. N.G.J., D.M.J., and J.P.A. were responsible for the experimental design. P.N.H. performed imaging and photobleaching experiments. C-L.C. performed 3D orbital tracking experiments. B.B. created fusion constructs.

## Data Availability Statement

Data will be made available to any interested parties

## Acknowledgments

Imaging and particle tracking were performed at the Laboratory for Fluorescence Dynamics (LFD) at the University of California, Irvine (UCI). The LFD is supported jointly by the National Institute of General Medical Sciences of the National Institutes of Health (P41GM103540), and UCI.

## References

1. Chappie, J. S.; Dyda, F., Building a fission machine--structural insights into dynamin assembly and activation. J Cell Sci 2013, 126 (Pt 13), 2773–84.

2. Morlot, S.; Roux, A., Mechanics of dynamin-mediated membrane fission. Annu Rev Biophys 2013, 42, 629–49.

3. Menon, M.; Schafer, D. A., Dynamin: expanding its scope to the cytoskeleton. Int Rev Cell Mol Biol 2013, 302, 187–219.

4. Sever, S.; Chang, J.; Gu, C., Dynamin rings: not just for fission. Traffic 2013, 14 (12), 1194–9.

5. Ferguson, S. M.; De Camilli, P., Dynamin, a membrane-remodelling GTPase. Nat Rev Mol Cell Biol 2012, 13 (2), 75–88.

6. Faelber, K.; Posor, Y.; Gao, S.; Held, M.; Roske, Y.; Schulze, D.; Haucke, V.; Noe, F.; Daumke, O., Crystal structure of nucleotide-free dynamin. Nature 2011, 477 (7366), 556–60.

7. Ford, M. G.; Jenni, S.; Nunnari, J., The crystal structure of dynamin. Nature 2011, 477 (7366), 561–6.

8. Srinivasan, S.; Dharmarajan, V.; Reed, D. K.; Griffin, P. R.; Schmid, S. L., Identification and function of conformational dynamics in the multidomain GTPase dynamin. EMBO J 2016, 35 (4), 443–57.

9. Zuchner, S.; Noureddine, M.; Kennerson, M.; Verhoeven, K.; Claeys, K.; De Jonghe, P.; Merory, J.; Oliveira, S. A.; Speer, M. C.; Stenger, J. E.; Walizada, G.; Zhu, D.; Pericak-Vance, M. A.; Nicholson, G.; Timmerman, V.; Vance, J. M., Mutations in the pleckstrin homology domain of dynamin 2 cause dominant intermediate Charcot-Marie-Tooth disease. Nat Genet 2005, 37 (3), 289–94.

10. Tanabe, K.; Takei, K., Dynamin 2 in Charcot-Marie-Tooth disease. Acta Med Okayama 2012, 66 (3), 183–90.

11. Bitoun, M.; Maugenre, S.; Jeannet, P. Y.; Lacene, E.; Ferrer, X.; Laforet, P.; Martin, J. J.; Laporte, J.; Lochmuller, H.; Beggs, A. H.; Fardeau, M.; Eymard, B.; Romero, N. B.; Guicheney, P., Mutations in dynamin 2 cause dominant centronuclear myopathy. Nat Genet 2005, 37 (11), 1207–9.

12. Wang, L.; Barylko, B.; Byers, C.; Ross, J. A.; Jameson, D. M.; Albanesi, J. P., Dynamin 2 mutants linked to centronuclear myopathies form abnormally stable polymers. J Biol Chem 2010, 285 (30), 22753–7.

13. Kenniston, J. A.; Lemmon, M. A., Dynamin GTPase regulation is altered by PH domain mutations found in centronuclear myopathy patients. EMBO J 2010, 29 (18), 3054–67.

14. Chin, Y. H.; Lee, A.; Kan, H. W.; Laiman, J.; Chuang, M. C.; Hsieh, S. T.; Liu, Y. W., Dynamin-2 mutations associated with centronuclear myopathy are hypermorphic and lead to T-tubule fragmentation. Hum Mol Genet 2015, 24 (19), 5542–54.

15. Massana Munoz, X.; Buono, S.; Koebel, P.; Laporte, J.; Cowling, B. S., Different in vivo impacts of dynamin 2 mutations implicated in Charcot-Marie-Tooth neuropathy or centronuclear myopathy. Hum Mol Genet 2019, 28 (24), 4067–4077.

16. Gomez-Oca, R.; Cowling, B. S.; Laporte, J., Common Pathogenic Mechanisms in Centronuclear and Myotubular Myopathies and Latest Treatment Advances. Int J Mol Sci 2021, 22 (21).

17. Tassin, T. C.; Barylko, B.; Hedde, P. N.; Chen, Y.; Binns, D. D.; James, N. G.; Mueller, J. D.; Jameson, D. M.; Taussig, R.; Albanesi, J. P., Gain-of-Function Properties of a Dynamin 2 Mutant Implicated in Charcot-Marie-Tooth Disease. Front Cell Neurosci 2021, 15, 745940.

18. James, N. G.; Digman, M. A.; Ross, J. A.; Barylko, B.; Wang, L.; Li, J.; Chen, Y.; Mueller, J. D.; Gratton, E.; Albanesi, J. P.; Jameson, D. M., A mutation associated with centronuclear myopathy enhances the size and stability of dynamin 2 complexes in cells. Biochim Biophys Acta 2014, 1840 (1), 315–21.

19. Chiu, C. L.; Digman, M. A.; Gratton, E., Measuring actin flow in 3D cell protrusions. Biophys J 2013, 105 (8), 1746–55.

20. Lanzano, L.; Gratton, E., Orbital Single Particle Tracking on a commercial confocal microscope using piezoelectric stage feedback. Methods Appl Fluoresc 2014, 2 (2).

21. Durieux, A. C.; Vignaud, A.; Prudhon, B.; Viou, M. T.; Beuvin, M.; Vassilopoulos, S.; Fraysse, B.; Ferry, A.; Laine, J.; Romero, N. B.; Guicheney, P.; Bitoun, M., A centronuclear myopathy-dynamin 2 mutation impairs skeletal muscle structure and function in mice. Hum Mol Genet 2010, 19 (24), 4820–36.

22. Gibbs, E. M.; Davidson, A. E.; Telfer, W. R.; Feldman, E. L.; Dowling, J. J., The myopathy-causing mutation DNM2-S619L leads to defective tubulation in vitro and in developing zebrafish. Dis Model Mech 2014, 7 (1), 157–61.

23. Zhao, M.; Smith, L.; Volpatti, J.; Fabian, L.; Dowling, J. J., Insights into wild-type dynamin 2 and the consequences of DNM2 mutations from transgenic zebrafish. Hum Mol Genet 2019, 28 (24), 4186–4196.

24. Cowling, B. S.; Chevremont, T.; Prokic, I.; Kretz, C.; Ferry, A.; Coirault, C.; Koutsopoulos, O.; Laugel, V.; Romero, N. B.; Laporte, J., Reducing dynamin 2 expression rescues X-linked centronuclear myopathy. J Clin Invest 2014, 124 (3), 1350–63.

25. Liu, N.; Bezprozvannaya, S.; Shelton, J. M.; Frisard, M. I.; Hulver, M. W.; McMillan, R. P.; Wu, Y.; Voelker, K. A.; Grange, R. W.; Richardson, J. A.; Bassel-Duby, R.; Olson, E. N., Mice lacking microRNA 133a develop dynamin 2-dependent centronuclear myopathy. J Clin Invest 2011, 121 (8), 3258–68.

26. Buono, S.; Ross, J. A.; Tasfaout, H.; Levy, Y.; Kretz, C.; Tayefeh, L.; Matson, J.; Guo, S.; Kessler, P.; Monia, B. P.; Bitoun, M.; Ochala, J.; Laporte, J.; Cowling, B. S., Reducing dynamin 2 (DNM2) rescues DNM2-related dominant centronuclear myopathy. Proc Natl Acad Sci U S A 2018, 115 (43), 11066–11071.

27. Gerber, D.; Ghidinelli, M.; Tinelli, E.; Somandin, C.; Gerber, J.; Pereira, J. A.; Ommer, A.; Figlia, G.; Miehe, M.; Nageli, L. G.; Suter, V.; Tadini, V.; Sidiropoulos, P. N.; Wessig, C.; Toyka, K. V.; Suter, U., Schwann cells, but not Oligodendrocytes, Depend Strictly on Dynamin 2 Function. Elife 2019, 8.

28. Achiriloaie, M.; Barylko, B.; Albanesi, J. P., Essential role of the dynamin pleckstrin homology domain in receptor-mediated endocytosis. Mol Cell Biol 1999, 19 (2), 1410–5.

29. Vallis, Y.; Wigge, P.; Marks, B.; Evans, P. R.; McMahon, H. T., Importance of the pleckstrin homology domain of dynamin in clathrin-mediated endocytosis. Curr Biol 1999, 9 (5), 257–60.

30. Lee, A.; Frank, D. W.; Marks, M. S.; Lemmon, M. A., Dominant-negative inhibition of receptor-mediated endocytosis by a dynamin-1 mutant with a defective pleckstrin homology domain. Curr Biol 1999, 9 (5), 261–4.

31. Tanabe, K.; Takei, K., Dynamic instability of microtubules requires dynamin 2 and is impaired in a Charcot-Marie-Tooth mutant. J Cell Biol 2009, 185 (6), 939–48.

32. Bitoun, M.; Durieux, A. C.; Prudhon, B.; Bevilacqua, J. A.; Herledan, A.; Sakanyan, V.; Urtizberea, A.; Cartier, L.; Romero, N. B.; Guicheney, P., Dynamin 2 mutations associated with human diseases impair clathrin-mediated receptor endocytosis. Hum Mutat 2009, 30 (10), 1419–27.

33. Liu, Y. W.; Lukiyanchuk, V.; Schmid, S. L., Common membrane trafficking defects of disease-associated dynamin 2 mutations. Traffic 2011, 12 (11), 1620–33.

34. Faelber, K.; Gao, S.; Held, M.; Posor, Y.; Haucke, V.; Noe, F.; Daumke, O., Oligomerization of dynamin superfamily proteins in health and disease. Prog Mol Biol Transl Sci 2013, 117, 411–43.

35. Muhlberg, A. B.; Warnock, D. E.; Schmid, S. L., Domain structure and intramolecular regulation of dynamin GTPase. EMBO J 1997, 16 (22), 6676–83.

36. Barylko, B.; Wang, L.; Binns, D. D.; Ross, J. A.; Tassin, T. C.; Collins, K. A.; Jameson, D. M.; Albanesi, J. P., The proline/arginine-rich domain is a major determinant of dynamin self-activation. Biochemistry 2010, 49 (50), 10592–4.

37. Kierdaszuk, B.; Berdynski, M.; Karolczak, J.; Redowicz, M. J.; Zekanowski, C.; Kaminska, A. M., A novel mutation in the DNM2 gene impairs dynamin 2 localization in skeletal muscle of a patient with late onset centronuclear myopathy. Neuromuscul Disord 2013, 23 (3), 219–28.

38. Fujise, K.; Okubo, M.; Abe, T.; Yamada, H.; Takei, K.; Nishino, I.; Takeda, T.; Noguchi, S., Imaging-based evaluation of pathogenicity by novel DNM2 variants associated with centronuclear myopathy. Hum Mutat 2021.

